# Preservation of Conditioned Behavior Based on UV Light Sensitivity in Dissected Tail Halves of Planarians- a Proof by DNN

**DOI:** 10.1101/2022.10.30.514395

**Authors:** Kensuke Shimojo, Eiko Shimojo, Reiya Katsuragi, Takuya Akashi, Shinsuke Shimojo

## Abstract

Planarians are aquatic worms with powerful regenerative and memory retention abilities. This paper examines whether a dissected tail half of a Planarian (Dugesia Dorotocephala) can retain and exhibit a previously-conditioned response, possibly before the regeneration of the head and the ganglia. We conditioned intact Planarians in a Pavlovian procedure with an electric shock (ES) as the unconditioned stimulus and weak ultraviolet (UV) light as the conditioned stimulus. Then, we dissected their bodies into halves, keeping the dissected tail halves. Starting from the 2nd day after dissection, we presented the same UV light 3 times daily while video-recording the responses. The recorded responses were then classified by a DNN: a VGG16 model was pre-trained by ImageNet for extracting features from images and additionally trained with 211 responses to ES and 118 to UV light before conditioning/dissection to categorize planarians’ reactions into “UV-induced” or “ES-induced” reactions. The cross-validated accuracy in categorization was 83.6%. We then let the DNN analyze 99 recorded responses to UV from 20 individual conditioned tail halves. 96.8 % of their reactions were classified as “ES-induced” (against 22.0% wrongly classified as “ES-induced” for unconditioned samples under UV), indicating they have shown the “Conditioned Response” (p<3.06E-30). This provides evidence that planarians can conserve and reveal a learned response even without the head/ganglia, as it takes approximately 7 days for the head/ganglia to regenerate versus the given 2-3 days. Although similar findings have been reported repeatedly in the literature, this is the first positive evidence with automated procedures and DNN classification. The result implies the presence of a decentralized nervous structure outside of its head/ganglia that allows a tail half to retain memory and execute motion accordingly, despite their cephalization.

## Introduction

Planarians, also known as flatworms, are a type of freshwater worms known for abilities of bidirectional regeneration and basic learning such as habituation ^1,2^. They have a cephalized nervous system, with a distinct head region that houses the ganglia (their brain-equivalent structure)^3^. They can also asexually reproduce by “fission” (splitting off a part of its tail), thus having the tail regenerate a new head and ganglia within a week^4^. However, for this way of reproduction to be effective, the tail must be able to survive in nature during the week of regeneration without its head, which houses many systems that play important roles in survival, such as the ganglia, the eyespots (visible light detection and aversion), and the auricles (food particle and touch detection). Indeed, the full regeneration of the ganglia requires up to 10 days^4^. As such, the tail half must be equipped with survival mechanisms that can function without its head, such as having ultraviolet light (UV) sensitive opsins distributed across the body. With this, planarian tail halves can detect/avoid sunlight to maintain moisture and stay hidden from predators^3^.

Another significant ability found in the tail is its ability to hold onto learned effects from habituation it experienced as a whole planarian ^2^.

This is a powerful tool for its survival as it offers higher familiarity in its given environment. However, it is a relatively simple form of learning compared to other learning types, such as Skinnerian or Pavlovian conditioning.

Such regeneration and survival abilities of the species have made it a perfect model to investigate the possibility of memory retention without the brain. Indeed, there are many classical studies employing conditioning^5^ procedures to indicate such memory retention beyond regeneration. The general procedure and logic go as such: the intact animals are trained to learn something, either in the conditioning or the habituation/addiction procedures. Then, they are dissected into halves, and the tail halves are tested for memory before the full regeneration of the neural ganglia.

Although there is ample classical evidence for such memory retention, these early studies classified the animals’ responses to the conditioned stimuli subjectively by the eyes, thus potentially vulnerable to bias and noise, providing no objective evidence. Also, most of the classical studies were not automated^8,9^, even though classical conditioning is known to be highly sensitive to timing, especially between the unconditioned and the conditioned stimuli. The current study aims to provide more objective evidence, employing a fully automated conditioning and testing protocol (by an Arduino microcontroller) and relying on a Deep Neural Network (DNN) to classify planarian behavior.

More specifically, the current study examines whether a dissected tail half of a planarian (Dugesia Dorotocephala) can retain learned behaviors from Pavlovian conditioning before its dissection to exhibit a previously conditioned response before the full regeneration of its ganglia. Pavlovian conditioning is beneficial to survival but is a relatively more complex and fundamentally different form of conditioning; studies on the cockroach showed that Skinnerian (operant) conditioning is possible without the central nervous system, but it is controversial if Pavlovian Conditioning is possible or not without the central nervous system^10,11^. It has been indicated that planarian tail halves can retain effects from Pavlovian conditioning and demonstrate their conditioned behavior before regenerating their head/ganglia^2^.

If our hypothesis is correct in that planarian tails can remember conditioned behavior for a better chance of survival and reproduction, we expect the dissected tail halves of the conditioned planarians to exhibit the conditioned response prior to the regeneration of the ganglia (<7 days after dissection). In this study, we put them under a Pavlovian Conditioning procedure with the following parameters: the conditioned stimulus is weak UV light, the unconditioned stimulus is an electric shock., and the response to the electric shock is a sharp contortion reaction. When this contortion response is triggered with only the conditioned stimulus (weak UV light), it would have become the conditioned response. Note that the intrinsic (pre-conditioning) response to this strength of UV light is very little and qualitatively different (*i*.*e*., stationary for the first several seconds, then slowly starts swimming) from that to the electric shock. Thus, we can train the DNN network with the responses of whole-body individuals to the electric shock and the weak UV light, such that it can classify a planarian’s reaction as either electric shock-like or UV-like. Then, we have it classify the tail halves’ responses to the UV light to see if the DNN mostly classifies them as responses to the electric shock, since this would imply that the tail halves showed the conditioned response and memory of conditioned behavior was retained.

## Results

After dissecting planarians that had been previously trained with Pavlovian conditioning, their dissected tails were able to exhibit their learned behavior prior to the regeneration of new ganglia.

We trained 20 individual planarians with a Pavlovian conditioning procedure such that they would start to associate exposure to weak UV with an electric shock (ES). As they were trained, they began reacting to the weak UV light with a contortion response similar to that toward the ES. (See the Methods for more details of the procedures.)

After the conditioning was successful, they were dissected, and the tail halves, on the 2^nd^ and 3^rd^ days after dissection (after a 1-day resting period), were exposed to the same weak UV to see if they exhibited the conditioned response (ES-like contortion reaction). A total of 99 responses from these 20 tail halves were recorded on video.

To objectively determine whether this recorded reaction is the conditioned response or not, we trained a deep neural network (DNN) to classify the recorded responses into “electric shock triggered responses” vs. “UV light triggered responses” (See the Methods section for more details). The DNN was trained with 211 control responses to ES and 118 to UV light with a separate, randomly chosen set of 20 whole-body planarians. Its cross-validated classification accuracy was 83.6%.

Then, this DNN categorized the 99 UV-stimulated responses from the tail halves recorded on days 2 and 3 after the dissection. 96.8% were classified as “ES-induced,” against 22.0% wrongly classified as “ES-induced” for unconditioned samples under UV, relative to 22.0 % as “ES-induced’ vs. % classified as “UV-induced” before the training (Fig. 1 shows the pooled data from days 2 and 3; see Supplementary Tables 1 and 2 online for original data).

**Fig 1.**
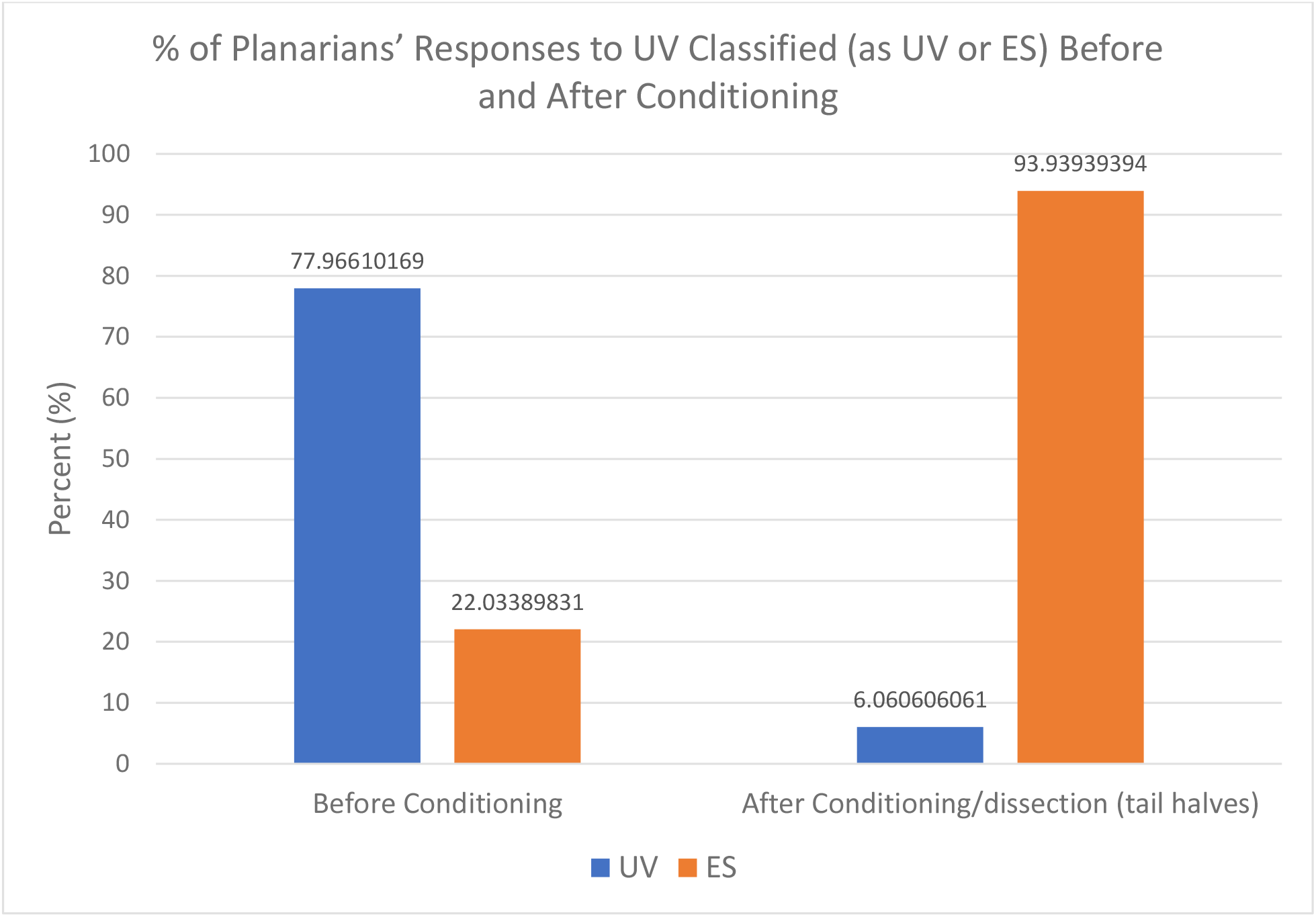
Classification of Responses to UV. % of Planarians’ Responses to UV Classified (as UV or ES) Before and After Conditioning. **x^2^=107.61, P=3.28E-25<.0000

Thus, as expected from the retention of the Pavlovian conditioning effect, the DNN classified a vast majority of the tails’ responses to UV as those to ES, relative to the latent 22% error on the DNN’s false ES categorization in the pre-conditioning responses. Statistical probability (p<3.28E-25<.00001) strongly suggests that the tail halves were able to retain information.

## Discussion

One of the biggest advantages of DNN-based classification is that it enables a classification that is purely based on objective features instead of subjective impressions by eye examination. As such, it opens a wide window to re-examine classical “findings” by eyes in animal behavior, neuroscience, and human psychophysics. In the current paper, we have applied a DNN-based classification algorithm (VGG16) to see if, after planarians have been conditioned and then dissected, their tail halves can reveal the learned behavioral responses even before regeneration of the neural ganglia.

The majority (>80%) of UV-induced responses on days 2 and 3 after the dissection (which was long before the 7th-10th day for the maturation of the ganglia) were classified by the DNN as “ES-induced,” confirming that the planarian can conserve and reveal a learned response without the head/ganglia. By employing the DNN and fully automated procedures (by the Arduino), we provided much more objective, convincing evidence than any in the classical planarian learning literature, essentially free from any bias and noise from eye examination. The results were consistent with the distributed nature of neural pathways and UV sensors in the animal^3^.

The result suggests the existence of a structure outside its ganglia that retains learned effects. This is consistent with the latest evidence on habituation, <u>but</u> it also shows a more sophisticated capacity of memory in this structure, as Pavlovian conditioning tends to be more complex than habituation. The results were also consistent with the distributed nature of neural pathways and UV sensors^3^.

Further, they were able to execute this function before the regeneration of their new ganglia, given the 1-2 day regeneration period they had before the tails were tested in our study vs. the 7-10 days needed for full regeneration.

This line of investigation is key to understanding the intricate interplay between the regeneration mechanism and the nervous system^12^. Along the line, it would be very beneficial to compare the biological mechanisms of regeneration and memory retention in the planarians with those in partly regenerative and non-regenerative species.

Since the entire body of the planarian is UV sensitive, it also raises a more drastic possibility of even learning without the ganglia (establishing new conditioning without the ganglia). As described earlier, Skinnerian conditioning is possible without the central nervous system in the cockroach, but this is not confirmed for Pavlovian conditioning^10,11^. It may also be worth seeing if the animal can reveal the learned behavior with the newly grown (not directly learned) ganglia.

### ProceduresThe overall procedures

can be summarized in the following three steps:

1. The selected animals (N=20) underwent the Pavlovian conditioning procedure, where the paired association of a neutral/conditioned stimulus with an unconditioned aversive stimulus/response was made. When successful, the conditioned stimulus alone can trigger the unconditioned response. Thus here, we employed an electric shock (ES) as the Unconditioned Stimulus (US) and weak UV light as the Conditioned Stimulus (CS). Their original, contortion-like response to the electric shock is considered the unconditioned response (UR). When this contortion response is observed toward the weak UV light, or the CS, it would be considered the Conditioned Response (CR), a learned behavior. In other words, the tail half showing a ES-like contortion response to only weak UV can be considered evidence for memory retention of the conditioned behavior.
2. The conditioned planarians are dissected. On the 2^nd^ and 3^rd^ days after the dissection, the tail halves are exposed to the same weak UV light, and their responses are recorded.
3. A DNN, pre-trained with not-yet-conditioned, whole-body Planarians’ responses to UV and ES, then categorizes the dissected tail halves’ responses to UV only as UV-like or ES-like responses. The majority of responses categorized as ES-like would then be taken as evidence for retained memory of conditioned behavior.

### Hardware Setup

The setup consists of a microcontroller (an Arduino), an overhanging projector camera, a UV flashlight weakened with UV filters, a 3D printed stand for the flashlight, a small aquarium with metal sheets (electrodes) attached to the sides, and a computer (See Supplementary Methods online for more details). The planarians go into the aquarium beneath the UV flashlight and the camera, such that the microcontroller controls the planarians’ exposure to UV and ES and starts/stops recording any of the planarians’ activities on the camera.

### DNN Setup and Training

We developed a video classification model for two planarian behaviors: the responses to ultraviolet light and electric shock. Many studies of video classification focused on a two-network architecture that extracts spatial features and temporal features from the video^13^. The two-stream models yield high-performance results in video classification. However, a large number of parameters are required in the model because the model fuses many layers of the neural network to extract spatiotemporal features from video, and the issue of overfitting can arise with a small dataset [2]^14^. Therefore, we developed a simple video classification model that expanded the scope of convolutional neural network (CNN) based image classification without complicated networks of the video classification model.

Our model is developed by expanding an image classification model with a CNN architecture such as VGG ^15^ and ResNet^16^. In our method, each frame of video is classified by the image classification model pre-trained by ImageNet^17^, and then, a mode value is calculated as a result of video classification with the image classification results. In our case, VGG16 is adopted as the backbone for image classification because it has the best performance. Moreover, we needed to modify our original videos of planarians recorded in our experimental environment and create a new dataset of videos as training data for the following 2 reasons: the model’s training is affected by the color difference between videos for planarians under UV vs. those under ES, and there are multiple planarians present in each video when the DNN needs to analyze one planarian’s motions at a time. To account for these issues, videos zoomed in on each planarian are extracted from a given video and are binarized (black/white). In training our video classification model, we used one frame per second of each one-planarian video, and all frames were input for analyzing the tails’ reactions.

### Experimental Procedure steps

1. Conditioning: repeated 180 conditioning trials (trial sequence shown below) on a batch of 20 planarians/day across 4 days:

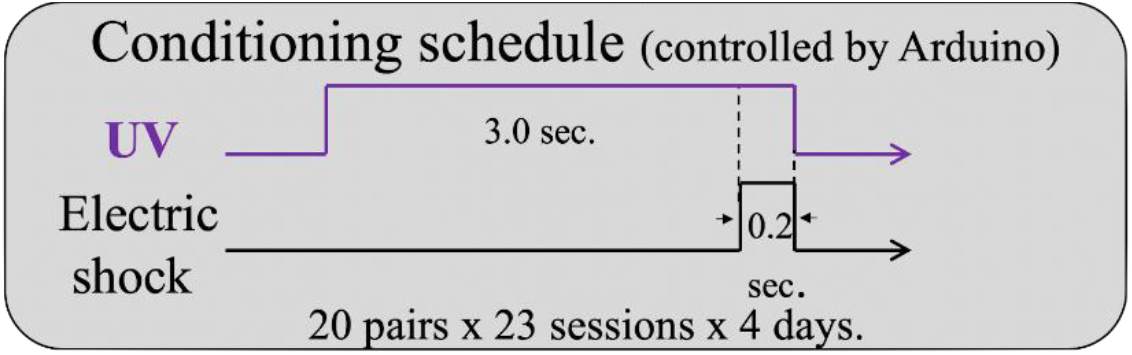 The following was done a total of 3 times for each of the 4 days at 8 am, 4 pm, and 12 am: 30 trials with 1-minute breaks in between, a 10-minute break, and another 30 trials with 1-minute breaks in between. We added the 1-minute breaks because, without them, the planarians became fatigued from the repeated electric shocks.
2. Exposed the 20 planarians under UV only and confirmed they have begun showing contraction response to weak UV alone (the CR): this indicates Pavlovian conditioning is established.
3. Manually dissected the conditioned whole-body animals with a scalpel, as close to the middle of the planarian as possible. The tail halves were given 1 full day to regenerate motor functions.
4. Regenerating tails were exposed to CS (weak UV) 3 times/day on the 2nd and 3rd days after dissection, with 3-minute breaks between each exposure. Their responses were recorded. Here, UV is turned on for 3 seconds, but recording time should last for an additional 7 seconds (a total of 10 seconds of footage), and a dim UV-free light should be on for the camera to capture the tail halves’ movements after UV exposure.
5. DNN Training: Recorded >100 control responses of a different group of 20 randomly chosen whole-body planarians to CS and US each. When recording responses to the US/electric shock, we used a lamp with a UV filter on or a UV-free lamp to light up the container, just enough for the planarians to be visible.
6. Recorded responses were fed to the DNN for training, such that it can categorize a given response as being more akin to the UV response or the ES response.
7. Recordings of tail halves’ reactions from step 4 were given to DNN for categorization

## Data Availability

Raw video data and DNN categorization results: https://drive.google.com/drive/folders/1Y47WyJePs8N9wF3ZW73_PGn03sJ5CLck?usp=share_link

## Acknowledgments

This project was supported by the Masason Foundation (Japan) and Chen Institute for Neuroscience at Caltech.

## Author Contributions

K.S. and S.S. contributed to conceiving and designing the experiment. E.S. and K.S. created the setup and ran the experiment. R.K. analyzed the data by training and running the DNN, and T.A. supervised this data analysis. K.S., E.S., S.S., and R.K. contributed to the preparation of the manuscript.

The authors declare no competing interests.

## References

1. Sánchez Alvarado A. Planarian regeneration: its end is its beginning. Cell. 2006 Jan 27;124(2):241–5. doi: 10.1016/j.cell.2006.01.012. PMID: 16439195.

2. McConnell JV, Jacobson AL, Kimble DP. The effects of regeneration upon retention of a conditioned response in the planarian. J Comp Physiol Psychol. 1959 Feb;52(1):1–5. doi: 10.1037/h0048028. PMID: 13641455. https://psycnet.apa.org/doi/10.1037/h0048028

3. Shettigar N, Joshi A, Dalmeida R, Gopalkrishna R, Chakravarthy A, Patnaik S, Mathew M, Palakodeti D, Gulyani A. Hierarchies in light sensing and dynamic interactions between ocular and extraocular sensory networks in a flatworm. Sci Adv. 2017 Jul 28;3(7):e1603025. doi: 10.1126/sciadv.1603025. PMID: 28782018; PMCID: PMC5533540.

4. Malinowski PT, Cochet-Escartin O, Kaj KJ, Ronan E, Groisman A, Diamond PH, Collins ES. Mechanics dictate where and how freshwater planarians fission. Proc Natl Acad Sci U S A. 2017 Oct 10;114(41):10888–10893. doi: 10.1073/pnas.1700762114. Epub 2017 Sep 25. PMID: 28973880; PMCID: PMC5642676.

5. Rescorla RA. Behavioral studies of Pavlovian conditioning. Annu Rev Neurosci. 1988;11:329–52. doi: 10.1146/annurev.ne.11.030188.001553. PMID: 3284445.

6. Shomrat, Tal and Michael Levin. “An automated training paradigm reveals long-term memory in planarians and its persistence through head regeneration.” Journal of Experimental Biology 216 (2013): 3799–3810.

7. Samuel, K., Suviseshamuthu, E.S. and Fichera, M.E. (2021) Addiction-Related Memory Transfer and Retention in Planaria. BioRxiv, doi: https://doi.org/10.1101/2021.09.12.459965.

8. Hicks, C., Sorocco, D. and Levin, M. (2006) Automated analysis of behavior: A computer-controlled system for drug screening and the investigation of learning. Journal of Neurobiology, 66-9, 977–990. https://doi.org/10.1002/neu.20290.

9. Blackiston, D., Shomrat, T., Cindy L. Nicolas, C.L., Christopher Granata, C. and Levin, M. (2010) A Second-Generation Device for Automated Training and Quantitative Behavior Analyses of Molecularly-Tractable Model Organisms. PlosOne, 6,(1): https://doi.org/10.1371/journal.pone.0014370.

10. Chen, W., Aranda, L., & Luco, J. (1970). Learning and long- and short-term memory in cockroaches. Animal Behaviour, 18, 725–732. https://doi.org/10.1016/0003-3472(70)90018-7

11. MacMillan, D. L. (1973). A classical conditioning paradigm for the study of learning in a ganglion of the cockroach (Periplaneta Americana). Animal Behaviour, 21(3), 492–500. https://doi.org/10.1016/S0003-3472(73)80009-0

12. Blackiston DJ, Shomrat T, Levin M (2015). The stability of memories during brain remodeling: A perspective. Commun Integr Biol 8: e1073424.https://doi.org/10.1101/2021.09.12.459965

13. Simonyan, K., & Zisserman, A. (2014). Two-stream convolutional networks for action recognition in videos. Advances in neural information processing systems, 27.

14. Jing, L., Parag, T., Wu, Z., Tian, Y., & Wang, H. (2021). Videossl: Semi-supervised learning for video classification. In Proceedings of the IEEE/CVF Winter Conference on Applications of Computer Vision (pp. 1110–1119).

15. Simonyan, K., & Zisserman, A. (2014). Very deep convolutional networks for large-scale image recognition. arXiv preprint 1409.1556.

16. He, K., Zhang, X., Ren, S., & Sun, J. (2016). Deep residual learning for image recognition. In Proceedings of the IEEE conference on computer vision and pattern recognition (pp. 770–778).

17. Russakovsky, O., Deng, J., Su, H., Krause, J., Satheesh, S., Ma, S., … & Fei-Fei, L. (2015). Imagenet large scale visual recognition challenge. International journal of computer vision, 115(3), 211–252.

