## Supplementary Table 1 for "Preservation of Conditioned Behavior Based on UV Light Sensitivity in Dissected Tail Halves of Planarians- a Proof by DNN"

### Supplementary Table 1: DNN Classification of Planarians' Responses to UV

| | UV | ES | Total |
|---|---|---|---|
| Before Conditioning | 92 (78.0%) | 26 (22.0%) | 118 (100%) |
| After Conditioning/dissection (tail halves) | 6 (6.04 %) | 93 (94.0%) | 99 (100%) |
| Total | 98 | 119 | 217 |
