## Supplementary Methods for "Preservation of Conditioned Behavior Based on UV Light Sensitivity in Dissected Tail Halves of Planarians- a Proof by DNN"

**Hardware setup:**

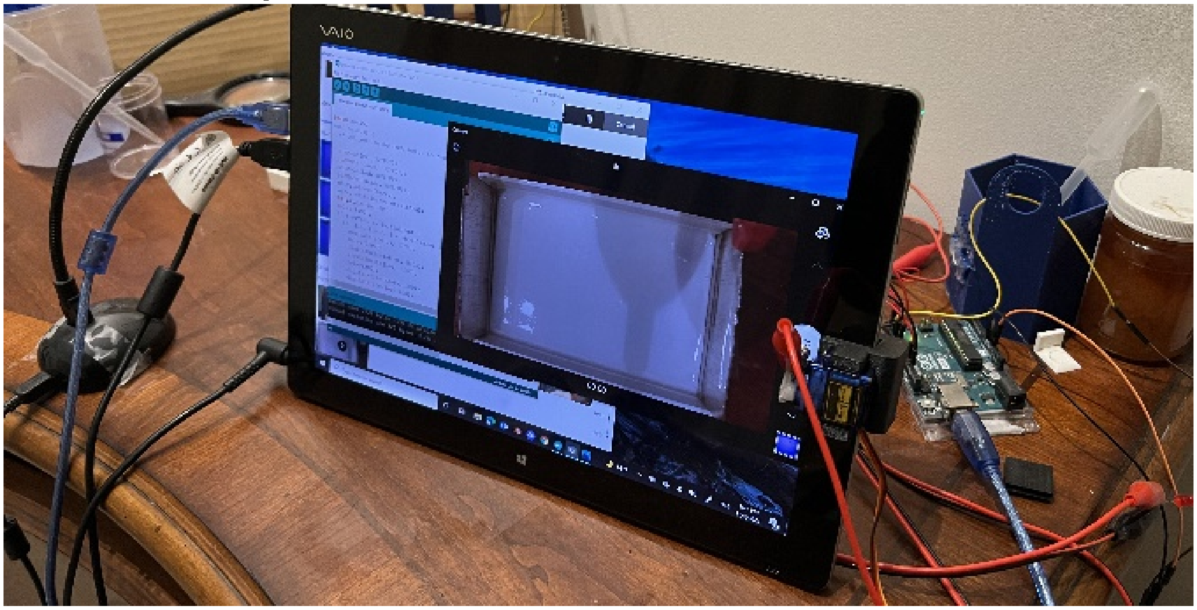

The setup consists of a microcontroller (such as an Arduino), an overhanging projector camera, a UV flashlight weakened with UV filters, a 3D printed stand for the flashlight, a plastic container with metal sheets hot glued on the sides, a tablet computer, and a servo motor with a metal tip attached to the tablet for pressing record.

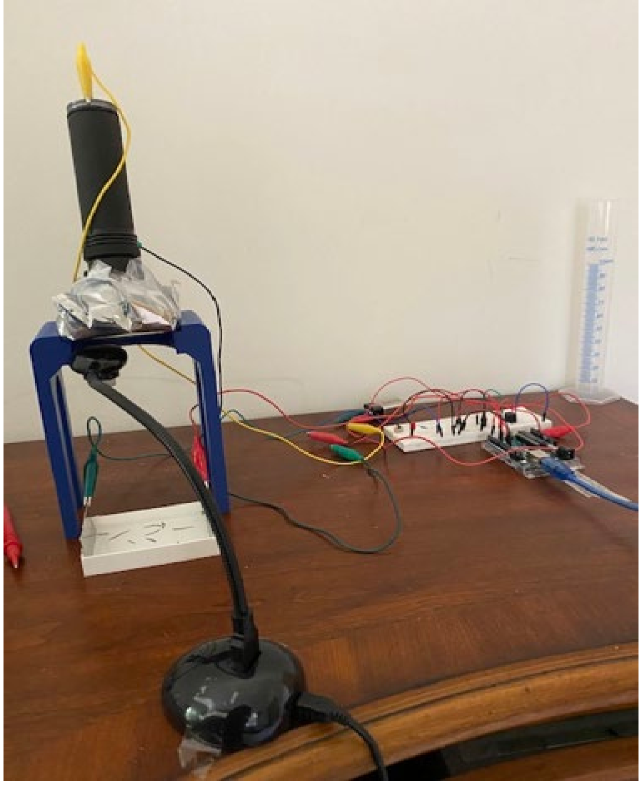

The tablet is connected to the camera as well as the Arduino, used to control the recording state and to run/modify the Arduino program. The Arduino uses transistors to control a 9V DC electric current through the testing aquarium using the 2 metal plates on the sides, as well as the UV light's on/off state and strength. It also controls a servo motor attached to the side of the tablet with an aluminum foil tip and an alligator clip such that it can interact with the tablet's touchscreen and control the camera's recording state.

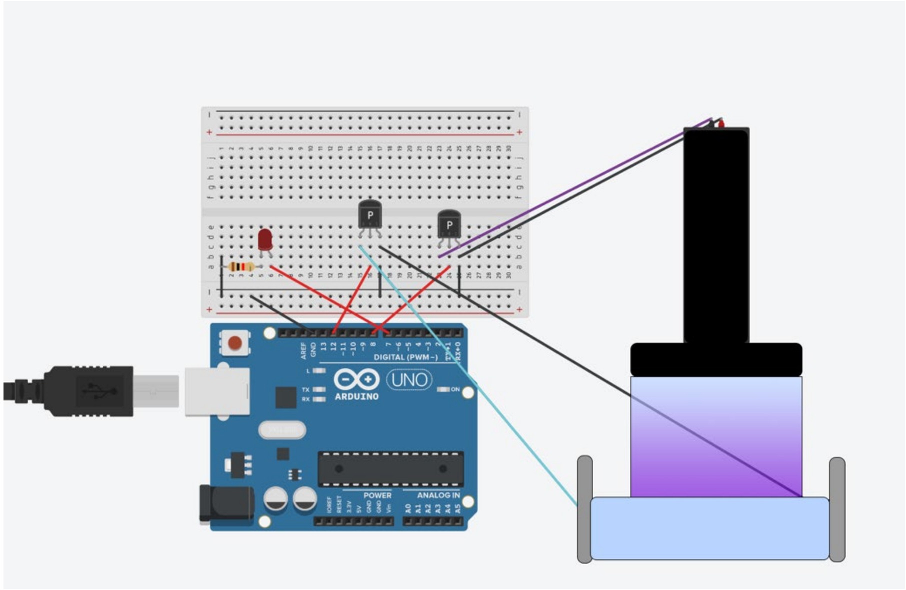

Above is a rough diagram of the wiring with the Arduino, with an extra LED as a sign of an electric shock being administered (not necessary).

Note that there should be a 9V voltage source (like a battery) in between the cyan and black wires connecting the transistor to the electrodes on the aquarium, and the servo motor for turning recording on/off is not shown.

The angled UV light holder, aquarium, and the servo motor holder for tapping record on tablet were modeled and 3D printed, although they can be substituted with any other similarly structured hardware.

**Determining the UV Strength:**
The UV light must be strong enough for the planarians to sense but weak enough to not cause a physical reaction. For this, we exposed the planarians to the UV light periodically, each time with decreasing strength (controlled either by attaching 50% UV filters or using the Arduino's PWM mechanism with the transistor) until the planarians no longer react to the UV light.
